# A meta-analysis of Neotropical butterflies reveals long-term signature of the Pebas wetland on Neotropical biogeography

**DOI:** 10.64898/2026.09.18.752375

**Authors:** Leidys Murillo-Ramos, André V.L. Freitas, Mar Repulles, Marianne Espeland, Pável Matos-Maraví, Nicolas Chazot

**Affiliations:** Department of Biology, Universidad de sucre, Sincelejo, Colombia; Laboratório de Ecologia e Sistemática de Borboletas, Departamento de Biologia Animal, Instituto de Biologia, Universidade Estadual de Campinas, Brazil; Biology Centre CAS, Institute of Entomology, České Budějovice, Czech Republic; Faculty of Science, University of South Bohemia, České Budějovice, Czech Republic; Leibniz Institute for the Analysis of Biodiversity Change (LIB), Museum Koenig, Bonn, Germany; Department of Ecology, Swedish University of Agricultural Sciences, Uppsala, Sweden; Department of Economic History, Lund University

**Keywords:** Neotropics, Pebas system, biogeography, diversification, meta-analysis, butterflies

## Abstract

The palaeogeographic history of the Neotropics has inspired multiple diversification models, emphasizing either the Andes or Amazonia as primary drivers of biodiversity. However, the extensive Pebas wetland system that covered western Amazonia during the Miocene has gained attention as a potential barrier to dispersal and diversification. This system may have contributed to the modern diversity imbalance between the Andes and Amazonia, known as the “Gentry pattern”. Most evidence for this hypothesis comes from isolated case studies and fossils. Here, we conduct a meta-analysis including 1,361 species of Neotropical butterflies to evaluate the role of the Pebas wetland on the dynamics of dispersal and diversification. We found that Andean- and Amazonian-centred clades show complementary but temporarily distinct biogeographic patterns. High Andean diversity reflects an Andean origin, with early occupancy and diversification occurring within the Andes, followed by post-Pebas dispersal into Amazonia. By contrast, Amazonian-centred clades, reflect an Amazonian origin, with an early diversification in Amazonia and subsequent post-Pebas colonization of the Andes. While several clades show little evidence of Pebas influence, our study suggests that the Pebas wetland did not uniformly restrict diversification, but rather selectively constrained dispersal between the Andes and Amazonia, preventing simultaneous diversification in both regions and thereby creating a diversity imbalance that characterizes the modern Neotropic.

## Introduction

The Andes and Amazonia together harbour the highest species richness in the Neotropic, yet the processes that generated their contrasting diversity patterns remain debated [1-4]. Narratives have generally focused either on Amazonia as a cradle of diversity or on the Andean uplift as a driver of allopatric and ecological speciation [5-9]. Amazonia has been portrayed as a stable, species-rich region where lineages originated and subsequently dispersed to other parts of the Neotropic, facilitated by its vast lowland forests and long-term climatic stability [7, 9, 10]. The long-debated Pleistocene Refugia Hypothesis [11] further emphasized recent contributions to Amazonian diversity as a result of repeated cycles of forest contraction and expansion during the last glacial periods.

In contrast, the uplift of the Andes created steep environmental gradients and novel habitats [2, 3, 12, 13], driving habitat specialization and ecological speciation [14, 15]. The Andes also created physical barriers that promoted geographic isolation, thereby increasing opportunities for allopatric speciation [12]. However, recent insights suggest a longer, more complex interplay between the Andes and the Amazonian basin, involving geological history, dispersal dynamics, and biogeographic barriers.

The uplift of the Andes played a key role in transforming the Amazonian landscape over time, particularly through the emergence and subsequent contraction of the Pebas System [4, 16]. The Pebas was a vast, dynamic wetland influenced by both fluvial and marine processes [4, 5, 17]. It originated around 23 Mya ago in the foreland basin east of the tropical Andes [4, 18] and expanded to cover, at its peak during the Middle Miocene, most of western Amazonia [17]. Aquatic conditions in the Pebas System were extended by large permanent lakes, and large areas supporting lowland swamp forests [4, 19].

Direct and indirect evidence suggest that the Pebas ecosystem may have strongly constrained the diversity of terrestrial fauna and flora in western Amazonia, before it was replaced by *terra-firme* and the establishment of modern rainforest [5, 20]. For example, during the Pebas period, the fossil record of terrestrial mammals reveals a significant decline in diversity before rebounding after the system disappeared [21]. Other case studies of plants, amphibians and butterflies have provided further evidence that the Pebas limited both dispersal and diversification in western Amazonia until its demise about 10 Mya [12, 15, 16, 18].

Antonelli and Sanmartín [22] extended this idea, proposing that the Pebas system exerted a lasting influence on Neotropical biogeographic patterns by creating an uneven distribution of diversity across the Neotropics. Their hypothesis built on Gentry [23]’s observation of two dominant diversity patterns in Neotropical plants, “Amazonian-centred” and “Andean-centred”, where species richness peaks in one of these centres relatively to the other [22]. According to Antonelli and Sanmartín, the Pebas system may have produced this dichotomy by inhibiting dispersal and diversification between eastern and western Amazonia.

More recently, Hoorn et al. [5] proposed a more nuanced view of the Pebas system as a “selectively permeable” biogeographic landscape. Rather than functioning solely as a constraint for dispersal, the Pebas wetlands may have allowed episodic and taxon-specific migration, particularly through intermittent fluvial connections between the Andes, Amazonian *terra-firme*, and the wetland itself. While this model reframes the Pebas as a dynamic and heterogeneous system whose permeability varied over time and among lineages [5], its link with modern geographic patterns, as suggested by Antonelli and Sanmartín [22], remains unclear.

One main obstacle in addressing this question is the limited number of studies explicitly testing for a Pebas scenario. Besides the general focus on the role of the Andean uplift and Amazonian rainforest, the evaluation of the Pebas scenario remains confined to isolated case studies with limited taxonomic scope and short evolutionary timescale [3].

Here, we unify these perspectives by asking whether differences in Pebas permeability among butterfly lineages explain Gentry-like richness patterns. We name as trans-Pebas distribution any lineage occupying a geographical area west of the Pebas system, mainly the Andean foothills and the Andes (but also Central-America), and cis-Pebas distribution as any lineage distributed mainly along the eastern margins of the Pebas system and eastern Amazonia. We derived three biogeographic scenarios that predict diversity patterns as follows:

1. Andean-centred diversity patterns emerged from an Oligocene - early Miocene trans-Pebas origin, low dispersal rates towards Amazonia during the Pebas period, and post-Pebas increase in Amazonian dispersal and diversification.
2. Amazonian-centred diversity patterns emerged from an Oligocene-Miocene cis-Pebas origin, low dispersal rate towards the Andes during the Pebas period, and post-Pebas increase in Andean dispersal and diversification
3. A balanced diversity pattern emerged within groups unaffected by the Pebas system, i.e. able to disperse and diversify simultaneously in cis- and trans-Pebas areas.

To test the adequacy of these scenarios beyond single case studies, we performed a meta-analysis of Neotropical radiations in the two most well-studied families of butterflies, the Nymphalidae and the Papilionidae. We compiled data for 15 Neotropical groups representing 1361 species, combined with phylogenetic framework, to perform ancestral range estimations under a single, homogeneous biogeographic model. We tested whether the geographical origin and the dynamics of dispersal and diversification conform to the scenarios described above and predict modern patterns of diversity.

## Material and Methods

### Clade selection, geographic and phylogenetic data

To move beyond case studies, we aimed to analyse multiple Neotropical radiations of butterflies within a comparative analytical framework that explicitly integrates Pebas permeability, diversification dynamics and directional dispersal across clades. These Neotropical groups were defined as clades whose diversification was centred on the Neotropics and clearly identifiable with a Neotropical common ancestor. We focused on two main families of butterflies for which comprehensive phylogenetic information exists: Nymphalidae and Papilionidae. Within Nymphalidae, we used the ancestral biogeographic estimation from Chazot et al. [24] to identify these Neotropical diversification events and extract time-calibrated trees. For Papilionidae, we used the family time-calibrated tree from Allio et al. [25]. We further refined the list of clades included in the analyses by setting a minimum of 10 species included in the tree. For each clade identified we extracted a summary tree and 100 trees randomly sampled from posterior distributions that varied in topology and branch length.

Each species included in the retained trees was assigned to the regions defined in our biogeographical model based primarily on a combination of previous publications, expert knowledge, and online resources such as GBIF [https://www.gbif.org], nic-funet [https://www.nic.funet.fi/] and iNaturalist [https://www.inaturalist.org/]. We defined seven biogeographic regions: Central America, Northern Andes, Central Andes, Southern Andes, Western Amazonia, Eastern Amazonia, Atlantic Forest. Any species occurring outside the Neotropics was simply assigned to an eighth region, “non-Neotropical”, without specifying further the distribution. Based on the taxa available in the trees, we then calculated species richness in Amazonia and in the Andes and quantified the relative difference in richness between the two regions. A clade was considered either Amazonian-centred or Andean-centred when the richness difference between regions exceeded more than 10% of the total species richness of the clade, regardless of whether one region contained most species. All trees not matching the condition above were assigned to a group with a “balanced diversity” pattern. This resulted in five Andean-centred clades, three Amazonian-centred clades and seven clades with balanced diversity (Supplemental S1).

### Ancestral biogeographic estimations

We performed ancestral state estimations using the C++ implementation of the DEC model, DECX [26]. Since we aimed at testing for an effect of the Pebas system, instead of *a priori* assuming it, we designed a non-stratified model, which only accounted for adjacency between regions. To account for phylogenetic and divergence-time uncertainty, we performed ancestral-state reconstructions and dispersal rate calculations across 100 posterior trees. For each dispersal event, we also resampled the timing of dispersal along the branch multiple times, as a uniform probability distribution. This approach incorporates variation in both tree topology, branch lengths and timing of events.

### Age and timing of local diversification

We estimated how long extant species have occupied either the Andes or Amazonia by recovering the timing of the oldest common ancestors that also occupied the same region without interruption. By estimating for each extant species its timing of origin in each region we aimed to (1) test if species richness variation between the Andes and Amazonia could result from a different amount of time each region was occupied (“time-for-speciation” hypothesis [14, 27]) and (2) estimate the proportion of lineages that emerged in each region before, during and after the Pebas period (“dispersal barrier” hypothesis). If the Pebas represented a barrier for dispersal and local diversification, higher species richness in a given region should be associated with longer occupation (and therefore species accumulation) in that region. Species found in the species-poor region should have originated more recently, after the Pebas retreat enabled dispersal. If the Pebas represented a constraint for dispersal and diversification in Andean and Amazonian centred clades, we also expected that the ancestral lineages in these two regions currently would have occupied the Andes and Amazonia either before or after but not during the Pebas when dispersal prevented the establishment of a new, distantly related lineages.

For each tip in the tree assigned to either the Andes or Amazonia, we traced back the oldest parental node with continuous occupancy and compared the age of Andean lineages with Amazonian lineages using violin plots. Additionally, we recorded every divergence time occurring along the path and ranked them from the oldest to the most recent, to visualize the timing of species accumulation in each region that resulted from these continuous local diversification events. These analyses were repeated on a posterior distribution of 100 trees.

### Dispersal rate through time

According to Antonelli and Sanmartin’s hypothesis, the Pebas played a significant role in isolating Amazonian lineages from Andean lineages. Under such a scenario, both cis-to trans-Pebas dispersal events and trans-to cis-Pebas dispersal events are expected to decrease during the Pebas period. To estimate the temporal variation in dispersal rate, we divided past evolutionary time into 2 million-year bins. Within each time bin, we calculated the number of dispersal events happening *towards* Amazonia and *towards* the Andes. Then, within each time-bin, we divided the sum of dispersal events by the sum of branch lengths to obtain the rate of dispersal. The DECX does not implement biogeographic stochastic mapping. Thus, to integrate in our analyses a distribution of probable ancestral state estimations we performed these analyses by sampling 1000 times a random timing of dispersal along the branch where a dispersal event happened and using the 100 ancestral state estimations performed on the posterior trees.

### Diversification rate

We compared the rate of diversification in Amazonia and in the Andes. If the Pebas acted as a barrier of dispersal, time-for-speciation and Pebas-constrained dispersal rates would be sufficient to generate asymmetrical biodiversity patterns. However, we also tested 1) whether variation in diversification rate between regions also predicted diversity patterns and 2) whether diversification rates changed over time in Amazonia and the Andes during and after the Pebas period.

We obtained branch-specific diversification rates using the implementation of ClaDS in Julia v.1.5.1 [28]. We fitted a ClaDS2 model with constant turnover on all trees from the posterior distributions of our dataset, accounting for missing taxa. We associated the estimated diversification rates and geographic distribution for each branch in the tree. Similar to dispersal rates, we divided time into 2 million-year bins. Within each time-bin, we multiplied each branch rate of diversification by its branch length, and finally, we averaged the rate of diversification from all lineages occupying either the Andes or Amazonia. To validate our results, we also compared CLaDS results with diversification rates estimated with BAMM over 100 million generations [29] (Supplemental S2).

## Results

When analysing the temporal origin of extant lineages occupying either Amazonia or the Andes, we identified two contrasting biogeographic patterns between Amazonian- and Andean-centred clades. In Andean-centred clades, the origin of lineages currently present in the Andes is concentrated around 20 Mya ago and to a lesser extent during the last 5 Mya with an important gap in between. In Andean-centred clades, Amazonian lineages mainly originated during the last 10 Mya (Figure 1A) but, to a lesser extent, also around 20-25 Mya ago. In Amazonian-centred clades, almost all Andean lineages appeared after the Pebas period (0-10 Mya, Figure 1B), whereas Amazonian lineages originated over a much longer time span of 30 Mya. Interestingly, there was no clear difference in the pattern of divergence time between Amazonian lineages and Andean lineages for clades of balanced diversity (Figure 1C), i.e. no obvious gap during the Pebas period.

**Figure 1.**
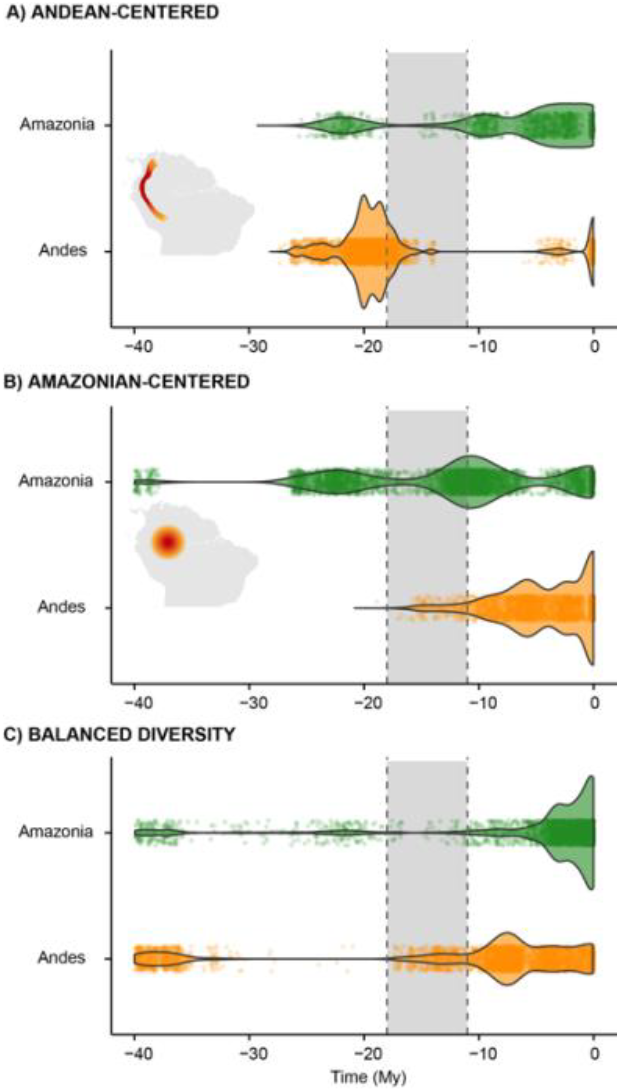
Distribution of ages of MRCA with continuous diversification in either the Andes or Amazonia to each tip of the trees for A) Andean-centred groups, B) Amazonian-centred groups and C) groups with balanced diversity. The figure shows the results for 100 posterior trees. The grey zone shows the peak of the Pebas period.

Lineage accumulation in both regions reflected the temporal patterns described above. In Andean-centred clades, species accumulated early in both the Andes and Amazonia (Figure 2A), but Amazonian lineages show a clear dampening of species accumulation during the Pebas period, while Andean diversity continues accumulating. In Amazonian-centred groups, Amazonian diversity started accumulated before the Pebas period, while species in the Andes accumulated almost entirely after the Pebas period (Figure 2B). Clades with balanced diversity show near identical dynamics of species accumulation in the Andes and Amazonia (Figure 2C), even though Andean lineages appear slightly more diverse than Amazonian lineages through history.

**Figure 2.**
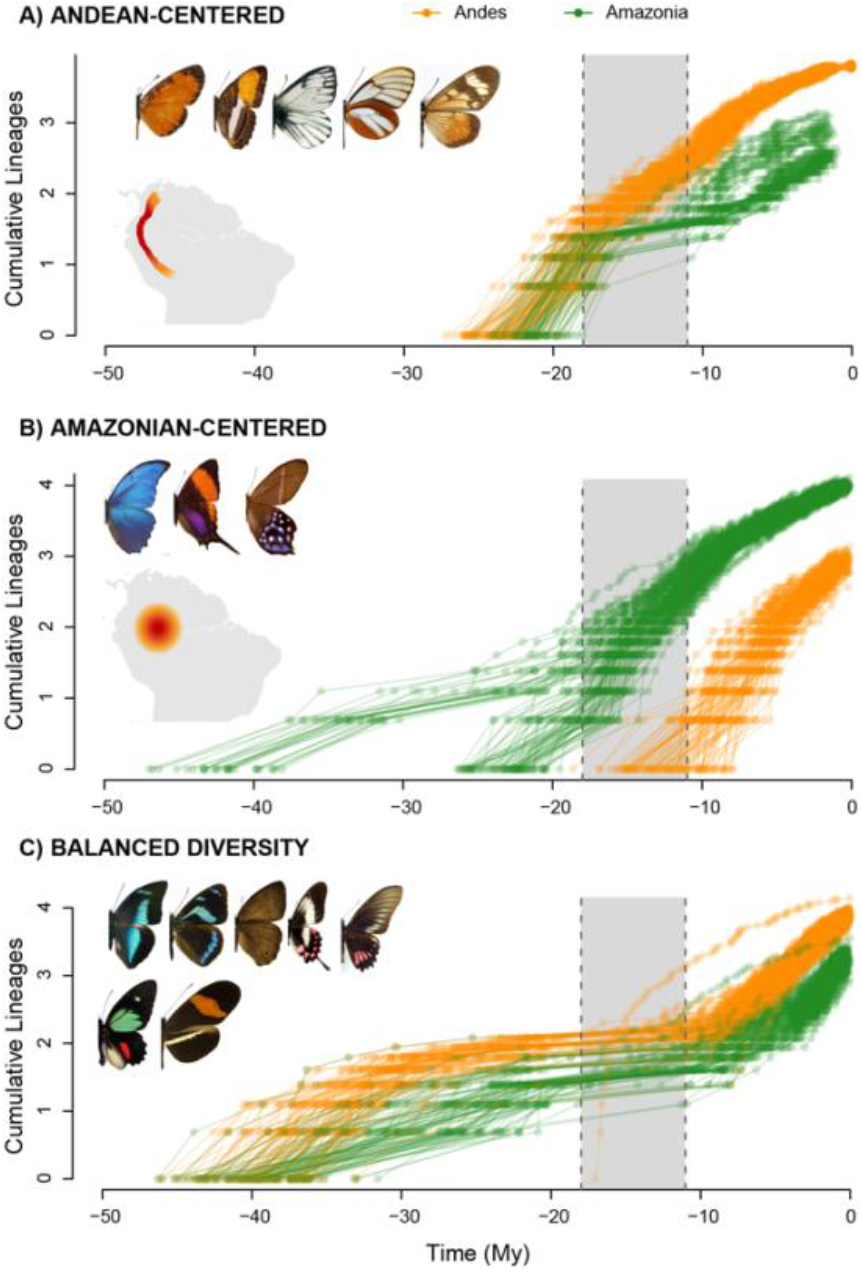
Accumulation of lineages in the Andes and Amazonia through time (in log-scale) for A) Andean-centred groups, B) Amazonian-centred groups, and C) groups with balanced diversity. The results are shown for 100 ancestral state estimations, each performed on a different posterior tree. The grey zone shows the peak of the Pebas period.

When analysing dispersal rates through time, we again found distinct patterns between our three categories (Figure 3). The Andean-centred groups showed early (pre-Pebas) dispersal in both Amazonia and the Andes followed by a dip in dispersal rate towards Amazonia during the Pebas period, especially towards Amazonia, before peaking again around 10 Mya followed by high dispersal rates during the second half of the Miocene (Figure 3A). Dispersal rate towards the Andes also showed a dip, delayed compared to dispersal towards Amazonia, rebounding during the last 5 Mya. In Amazonian-centred clades, dispersal towards the Andes remained extremely low until peaking during the last 10 Mya ago. Dispersal towards Amazonia, instead, remained high pre- and during the Pebas period but declining after the Pebas retreat (Figure 3B). In clades with balanced diversity, we found relatively constant dispersal rates towards Amazonia over more than 30 Mya of evolution. Dispersal in the Andes increased during the early Miocene, reaching a plateau during the early Pebas period at a similar rate with Amazonian dispersal (Figure 3C). Importantly, there was no variation associated with the Pebas or post-Pebas period.

**Figure 3.**
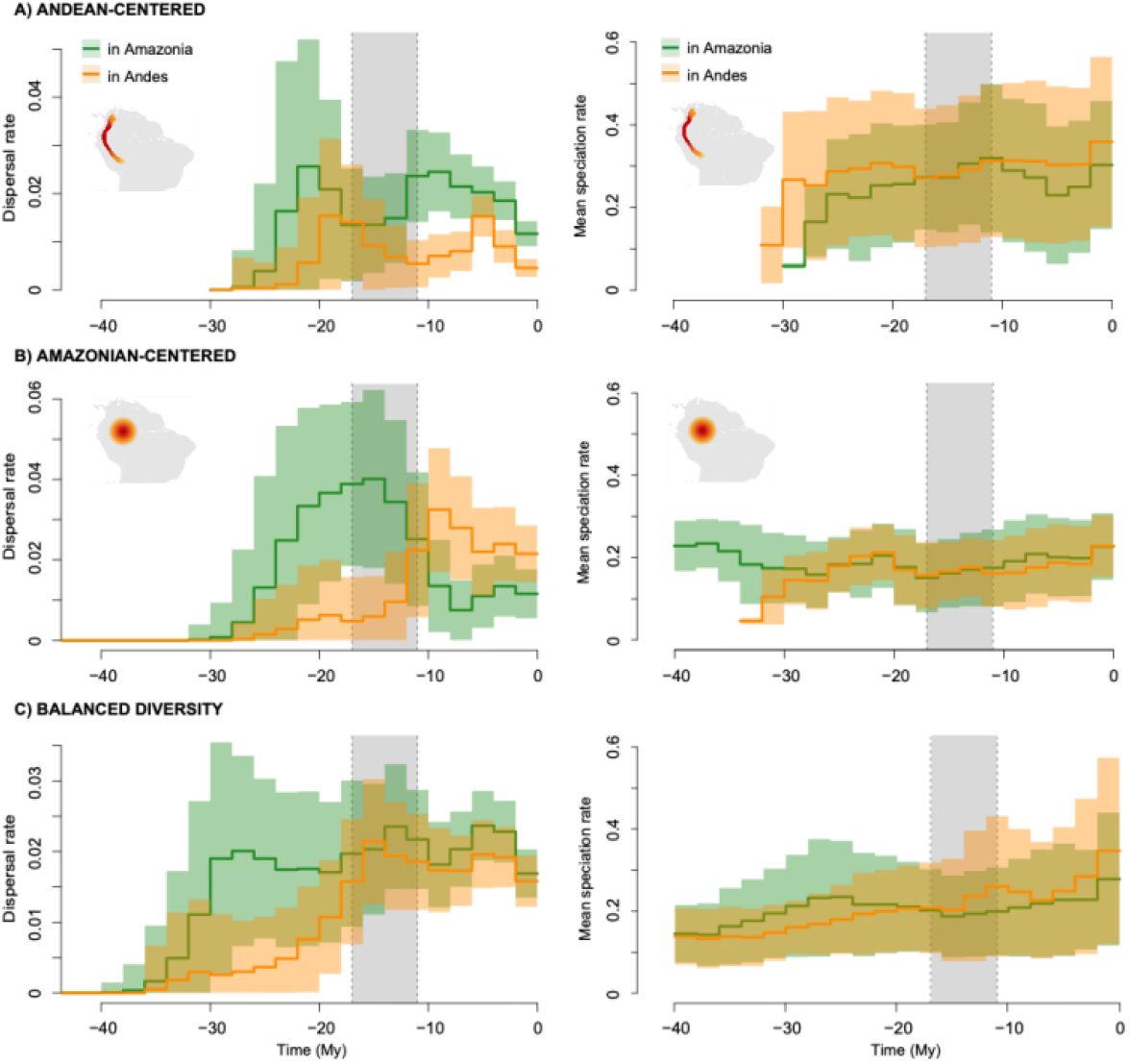
Dispersal rates (left panels) and diversification rates (right panels) through time in the Andes and Amazonia for A) Andean-centred groups, B) Amazonian-centred groups, and C) groups with balanced diversity. The shaded areas show the 95% interval of dispersal and diversification rate calculated after performing 100 ancestral state estimations, each performed on a different posterior tree and sampling 1000 times dispersal events along the branches in the tree. Bold lines indicate the mean rate for each time-bin. The grey zone shows the peak of the Pebas period.

The comparison of speciation rates showed no clear difference between regions and clades (Figure 3, 4). Through time, the speciation rate either remained stable over time or generally slowly increased towards the present in both regions. Within Andean-centred groups and within Amazonian-centred groups, there was no variation in speciation rate between Andean and Amazonian lineages that could explain the imbalance in species richness. Andean-centred clades appear to have generally higher speciation than Amazonian-centred or balanced diversity clades, whether they occupy the Andes or Amazonia. Thus, contrary to all previous results, we found no general support for contrasting patterns between Andean- and Amazonian-centred clades that would explain the imbalance in species richness.

**Figure 4.**
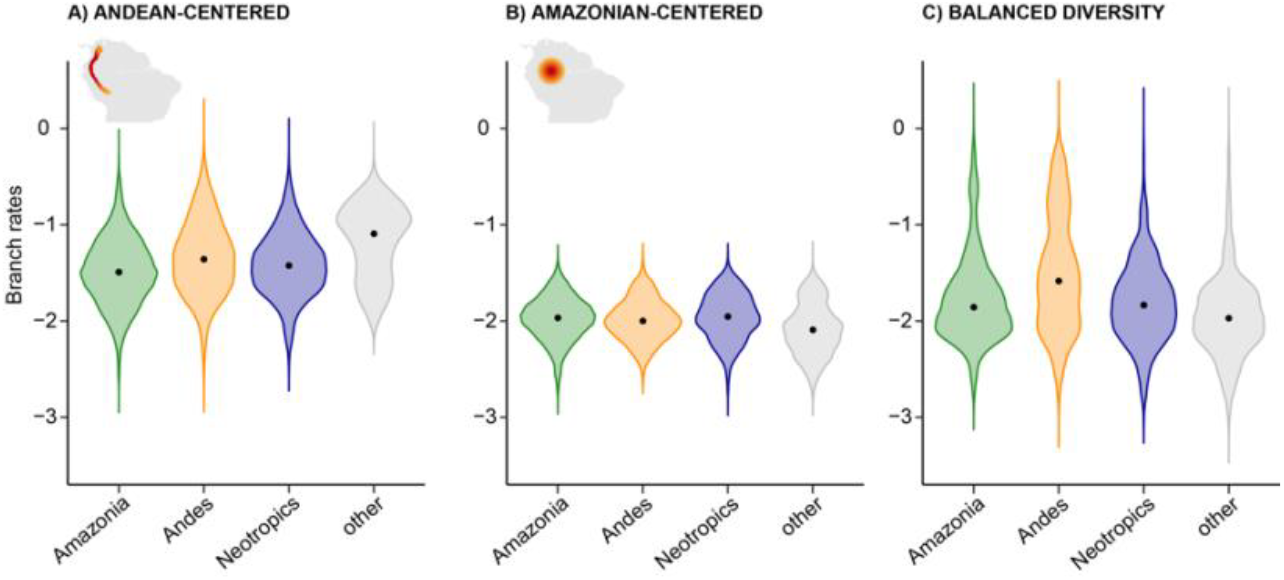
Distribution of branch-specific rates of speciation from CladS for A) Andean-centred groups, B) Amazonian-centred groups, and C) groups with balanced diversity. “Amazonia” refers to branches inferred as Amazonian, “Andes” refers to branches inferred as Andean, the “Neotropics” refers to branches inferred as all South-American areas except Amazonia and Andes, “other” refers to branches inferred outside the Neotropics.

## Discussion

In this study, we aimed at testing the role of the Pebas system in generating modern patterns of species richness in the Neotropics. We found that groups of butterflies that exhibit a Gentry-like pattern – either Andean-centred or Amazonian-centred species richness – conform particularly well with the model of dispersal and diversification constrained by the Pebas [22].

### Support for the Pebas Barrier Hypothesis

Our results show an early, pre-Pebas occupation of both the Andes and Amazonia by Andean-centred groups, but a clear divergence happening during the Pebas period. Most species nowadays present in Amazonia within these clades trace their Amazonian origin to the last 10 Mya or, to a lesser extent before the Pebas, but almost never during the Pebas period itself (Figure 1). This is consistent with the dip in dispersal rate and lineage accumulation in Amazonia and the following increase in Amazonian dispersal around 10 Mya and during the late Miocene (Figure 3). This pattern is in line with our expectations under a scenario where the Pebas acted as a constraint for eastward dispersal during the early Miocene (23–10 Mya), followed by a post-Pebas expansion and diversification in Amazonia [3, 4]. In contrast, Amazonian-centred groups started diversifying in Amazonia long before the Andes, probably in Eastern Amazonia unoccupied by the Pebas or *terra-firme* areas at the eastern margin of the Pebas ecosystem. These lineages clearly failed to disperse and diversify towards the Andes before the demise of the Pebas system, around 10 Mya, when we inferred a clear peak of dispersal events into the Andes (Figure 3). Consistent with this scenario, nearly all Andean species of the Amazonia-centred group trace their origin after the Pebas (Figure 1). These temporal patterns are again consistent with the predictions from Antonelli & Sanmartín’s barrier model, where ecological conditions associated with the Pebas, constrained early colonization of trans-Pebas regions. Our results also suggest an association between the centre of diversity and the centre of origin, especially for Amazonian-centred groups in line with the more general time-for-speciation scenario [14, 27], where longer residence time predicts higher local species richness.

Our results are consistent with previous studies on amphibians [30, 31] and wasps [32], which also suggest that the Pebas system functioned as a barrier to diversification in western Amazonia during the Miocene. Amphibians such as the genus *Allobates* show a pattern of early Miocene lineage accumulation in regions affected by the Pebas wetlands, followed by reduced diversification as the system expanded and fragmented habitats [20, 30, 31]. Similarly, Epiponini wasps underwent a rapid burst of diversification, with a subsequent slowdown coinciding with the ecological constraints imposed by the Pebas system [[32].

Evidence from other taxa suggests that *in situ* diversification in western Amazonia was more likely to have occurred during the final stages of the Pebas system [30, 33-35]. After the contraction of the Pebas, the emergence of new *terra-firme* led to a reshaped riverine network and increased availability and heterogeneity of terrestrial habitats [1, 3]. This environmental transformation likely facilitated the subsequent diversification and expansion of terrestrial organisms throughout Amazonia [12, 35]. Future meta-analyses designed to test the same predictions for other groups outside butterflies are needed to assess the generality of our findings.

### Semi-permeable Pebas Landscape

Clades with a balanced diversity seem to be less affected by the presence of the Pebas system. We did not find an obvious variation in dispersal rate during or after the Pebas system. The origin of extant Andean and Amazonian lineages can be traced back to the Andes and Amazonia respectively before, during and after the Pebas period, without any clear Pebas-associated variation. These findings suggest that ecological connectivity existed between the Andes and the western Amazonia even during the Pebas period, supporting the premise that the Pebas functioned as a permeable biogeographic system for certain taxa [1, 5], providing dispersal and local diversification opportunities. This pattern aligns with studies on plants [19, 36], which have indicated that environmental conditions along the margins of the Pebas region favoured the vegetation that eventually transitioned and adapted to *terra-firme* habitats [19, 35].

Although the mechanisms driving speciation are diverse, the recurrent movement of butterfly lineages between the Andes and Amazonia may reflect the constraints of adult butterfly ecology, particularly the need to locate suitable host plants, habitats, and mates [37-39]. While caterpillars are often restricted by the distribution of their host plants, adults can disperse over broader distances [37, 40, 41]. The observed dispersal and diversification of butterflies may reflect parallel responses to the biogeographic history and environmental barrier of their host plant groups [42][41], which are themselves constrained by the Pebas system. Other ecological traits, such as long-term dispersal capacity or microhabitat specialization, could also play a critical role in shaping lineage movements [43] and responses to historical barriers like the Pebas system. Lineages could probably occupy ecological zones that were not submerged, or simply had traits that allowed them to survive or skirt the Pebas system [5]. These traits, like highly mobile capacity, generalist diet or canopy adaptations might help explain why some groups appear less affected by landscape changes than others [44-46]. Similarly, species with broad environmental tolerance could have found more distinct niches in the Amazonia, leading to adaptive radiation. While these traits offer plausible explanations for why certain clades show balanced diversity, our current knowledge of the relevant ecological and life-history traits in these butterflies remains limited, preventing us from rigorously testing such hypotheses at this stage.

### Diversification rates: limited impact

We did not detect major differences in diversification rates between regions within each group. Although both the Andes and Amazonia show increasing rates of speciation throughout the Miocene, these rates do not differ significantly between the two regions. This suggests that, for the butterfly groups analysed here, time and dispersal opportunity—rather than intrinsic differences in regional speciation rates—are the primary drivers of present-day richness imbalance [47, 48].

This suggests that richness imbalance may not be the result of where species diversify, but when diversification opportunities arose and how lineages moved across biogeographical regions [47-49]. Amazonia and the Andes have both experienced major geographical and ecological changes since the Mid-Eocene, but if diversification rates remain broadly similar across regions, other extrinsic forces rather than diversification capacity have triggered present-day diversity, leading to relatively stable rates across different environments. However, we cannot discard that limited taxon sampling, the small tree sizes and incomplete phylogenies reduce the power to detect rate heterogeneity across regions in our study.

The direct or indirect consequences of Andean and Amazonian dynamics are particularly complex to decipher; however, by comparing our results to the existing biogeographical studies, we open possibilities for an ongoing revaluation of how biogeographical history shapes macroevolutionary patterns across the Neotropics. It remains essential to assess the extent to which our conclusions apply beyond butterflies and to other terrestrial taxa. While the uplift of the Andes has long been emphasized as a major driver of diversification, our findings suggest that the historical dynamics of the Pebas system may have played an equally significant role, substantiating Antonelli and Sanmartín [22] original hypothesis.

## Conclusions

Our findings reinforce the idea that large-scale palaeoenvironmental changes can leave enduring biogeographic signatures. On the one hand, the Andean uplift created the conditions for repeated diversification across altitudes and cordilleras, contributing significantly to Neotropical diversity. On the other hand, we show that Miocene flooding events had a strong impact on biotic interchanges, creating a strong imbalance in diversification between regions and resulting in the modern Gentry-pattern. Applying the framework presented in this study to other groups is now needed to assess the generality of our findings beyond butterflies.

## Notes

### Competing Interest Statement

The authors have declared no competing interest.

### Funding

- The Swedish Research Council for Environment, Agricultural Sciences and Spatial Planning: Grant number 2023-00339
- Conselho Nacional de Desenvolvimento Cientí-fico e Tecnológico – CNPq: grants 304291/2020-0, 408764/2024-4 and 317331/2025-6
- Fundação de Amparo à Pesquisa do Estado de São Paulo: FAPESP, grants 2021/03868-8 and 2023/04393-9
- Bilateral Czech Science Foundation grant: GACR 22-35084J
- Vetenskapsrådet: Grant no: 2023-04948

